# Size Selection of Giant Unilamellar Vesicles (GUVs) via Modified cDICE Method

**DOI:** 10.1101/2024.10.23.619772

**Authors:** Ariel Chen, Shachar Gat, Lior Ohana, Evgeni Yekimov, Yoav Tsori, Anne Bernheim-Groswasser

**Author notes:** **Corresponding Author Anne Bernheim-Groswasser**-Department of Chemical Engineering and the Ilse Kats Institute for Nanoscale Science and Technology, Ben-Gurion University of the Negev, Beer-Sheva, Israel.

## Abstract

The production of giant unilamellar vesicles (GUVs) plays a pivotal role in various scientific disciplines, particularly in the development of synthetic cells. While numerous methods exist for GUV preparation, the modified continuous droplet interface crossing encapsulation (cDICE) method offers the advantages of simplicity and high encapsulation efficiency. However, a significant limitation of this technique is the generation of vesicles with a broad size distribution and the inability to control the desired size range. This raises a key question: Can the modified cDICE method be optimized to produce GUVs with controlled size distribution? In this study, we examined the effects of two experimental parameters—rotation time (*t*_ROT_) and the angular frequency (*ω*) of the cDICE chamber—on the size distribution of GUVs. Our results show that reducing either the angular frequency or rotation time shifts the size distribution toward larger vesicles, enabling effective size selection. These findings are further supported by a physical model, which provides insights into the mechanisms underlying size selection. This work demonstrates that control over GUV size distribution can be achieved through straightforward adjustments of system parameters. The ability to fine-tune vesicle size offers researchers a powerful tool for developing customizable experimental systems for synthetic biology and related fields.

---

Synthetic cells are invaluable tools for exploring the complexities of cellular functions and advancing the design of novel biotechnologies. By constructing artificial cells, scientists can investigate fundamental biological processes in a simplified and controlled environment, leading to new insights and innovations. Various approaches exist for producing cell-mimicking compartments, including water in oil droplets, polymersomes, and liposomes.

Giant unilamellar vesicles (GUVs) stand out for several reasons. Like living cells, GUVs are large spherical compartments enclosed by a lipid bilayer. Their cell-sized dimensions (5-50*µ*m) that surpass the optical resolution limit facilitate their investigation using light microscopy. Their structural similarity to real cells, in terms of size and lipid composition, coupled with their customized nature, makes GUVs ideal for using them for probing cellular processes under controlled conditions and for developing bio-inspired applications in medicine, diagnostics, and synthetic biology.

GUVs serve as a versatile model system for characterizing the mechanical properties of lipid membranes, with or without embedded membrane proteins. ^1–4^ GUVs are also useful for studying the impact of membrane-bound proteins and membrane curvature on the self-organization of cytoskeletal networks, ^5,6^ and the effect of integrin-mediated surface adhesion on cell spreading. ^7^ Additionally, vesicles serve as bioreactors, ^8^ which provides a controlled environment for biochemical reactions and can also be utilized in drug delivery applications. ^9^

GUVs can be formed by different methods and technologies (e.g., electroformation, ^10^ gel-assisted swelling, ^11^ inverted emulsion, ^12^ and microfluidic-assisted platforms. ^13,14^ These techniques differ in the yield and encapsulation efficiencies, as well as in the size range and polydispersity of the generated GUVs. Some techniques are limited to using lipid composition with a low proportion of charged lipids and solutions with relatively low ionic strength, such as the electroformation technique. In particular, using an electric field limits the use of sensitive biological materials, notably required in developing a synthetic cell. Many of these methods also entail a time-consuming process to generate the vesicles, limiting their use for observing dynamic transient states.

The continuous droplet interface crossing encapsulation (cDICE) method offers several advantages over these techniques. It is a rapid and inexpensive method, demonstrating a high GUVs’ production yield and elevated encapsulation efficiency. ^15^ Furthermore, it can work with lipid compositions containing high proportions of charged lipids and with high ionic strength solutions like those present in the cellular environment. ^16^

The cDICE method consists of three main steps. The first includes the preparation of a lipid-in-oil mixture (LOM) with mass density *ρ*_LOM_ and dynamic viscosity *µ*_LOM_ (Fig. 1) through a process called “solvent-shifting” or “Ouzo Effect”. ^17^ In this step, lipids are first dissolved in a good solvent (e.g., chloroform) and then mixed with a non-miscible oil phase (e.g., a mineral and silicon oil mixture ^15^), which triggers the formation of lipid aggregates (blue and red spots in schematic diagram/Image (i) in Fig. 1). Then, an aqueous phase (denoted the ‘outer solution,’ O) and the LOM, are dripped in sequential order into a chamber rotating at an angular frequency *ω* (Fig. 1). The applied centrifugal forces trigger their phase separation, with the ‘O’ phase being pushed toward the outer chamber edge owing to its higher mass density, *ρ*_O_, and the establishment of a lipid monolayer at their interface. In the third step, aqueous droplets of the solution (denoted as the ‘inner solution’ I) are dripped into the rotating chamber. Setting the mass density of the different solutions to *ρ*_I_ *> ρ*_O_ *> ρ*_LOM_, ensures that the aqueous droplets are driven towards the LOM/O interface, where they undergo membrane zipping, and are eventually released in the ‘O’ phase as GUVs (see image (ii) in Fig. 1).

**Figure 1:**
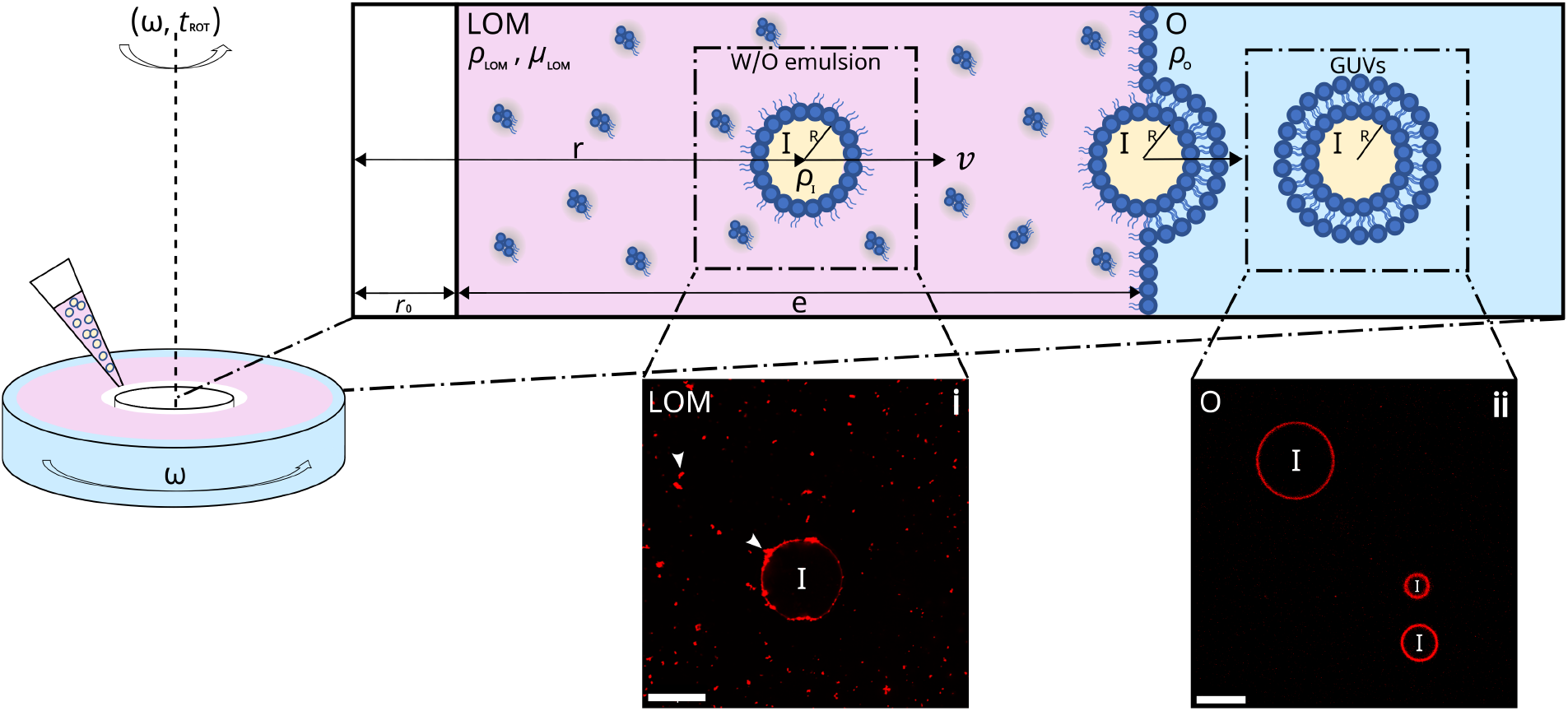
GUVs formation using the modified cDICE method - Schematic diagram. An outer aqueous phase (‘O’) of mass density *ρ*_O_ and a lipid-in-oil mixture (LOM, lipids are marked in blue) of mass density *ρ*_LOM_ and dynamic viscosity *µ*_LOM_ are sequentially dripped into a rotating chamber operating at an angular frequency *ω*. The centrifugal force generated during rotation induces phase separation and promotes the formation of a lipid monolayer at their interface. Subsequently, water-in-oil emulsion droplets of sample solution (‘I’) and mass density *ρ*_I_ are introduced in the rotating chamber. These droplets migrate at a velocity *v*, which depends on their radius *R* toward the LOM/O interface. There, they undergo membrane zipping and are released as GUVs in the outer aqueous ‘O’ solution. Only droplets that reach the LOM/O interface at *r*_0_ + *e* in a time shorter than the overall experiment time (i.e., the rotation time *t*_ROT_) end up as GUVs. Also depicted are confocal images of: i) a water-in-oil droplet in the LOM mixture. The red spots are lipid aggregates (white arrow heads) present in the LOM mixture and the W/O droplet interface. ii) GUVs immersed in the outer ‘O’ phase. Membrane composition (molar percentage): 99.89% DOPC, 0.10% Liss Rhodamine-PE, 0.01% DSPE-PEG(2000)-Biotin. Scale bars are 20*µ*m.

Droplets can be introduced bare (i.e., uncoated) using a glass capillary (as per the original cDICE method ^16,18,19^) or pre-coated with a lipid monolayer as a water-in-oil (W/O) emulsion (as per the modified cDICE method ^15^). Fig. 1 (Image (i)) depicts an example of such an emulsion droplet. The use of a preformed emulsion is greatly advantageous over the use of naked droplets. First, this method is technically less challenging and does not require expensive equipment beyond standard laboratory equipment, such as a pipette puller. Moreover, an essential factor to consider when using naked droplets is the ‘time of flight,’ which is the duration it takes for the droplets to travel a distance *e*, corresponding to the LOM layer thickness, and reach the LOM/O interface. This time should be sufficiently long to ensure that all naked droplets, regardless of their radius *R*, reach the LOM/O interface fully saturated with lipids, a prerequisite for their eventual development into a GUV. ^18^ The use of an emulsion resolves this issue. We note, however, that a significant limitation of the modified cDICE method is its difficulty in consistently controlling the size of the GUVs produced, ^15^ which primarily depends on the distribution of droplets’ size introduced in the chamber.

Here, we demonstrate efficient control of the generated GUV size and size distribution by using the rotating chamber as a size sorting or selecting element. We demonstrate how such a size selection can be achieved through proper adjustments of the rotation time (*t*_ROT_) and angular frequency (*ω*). Our experimental results are supported by a physical model, which allows us to better understand how size selection is achieved and what the underlying mechanism governs this process.

## RESULTS AND DISCUSSION

### In Situ Size Selection of GUVs

#### Effect of Rotation Time on GUVs’ Size Distribution

GUVs were generated using the modified cDICE method. ^15^ We conducted a set of experiments at a fixed rotation speed *ω* = 1600 rpm to explore the effect of the rotation time *t*_ROT_ on GUV size distribution (Fig. 2). Each experiment includes the following time steps from which we calculate the rotation time: (i) the emulsion dripping time = 40 sec, constant in all studied systems, independently of the rotation speed, (ii) the time the chamber rotates steadily at *ω* = 1600 – four different times were tested 15, 45, 180, and 1200 sec, and (iii) the time it takes for the chamber to decelerate from *ω* = 1600 to 0 rpm at a rate of 100 cycles/sec (= 16 sec). In Fig. 2, we show the results obtained for the corresponding four rotation times: 71, 101, 236, and 1256 sec.

**Figure 2:**
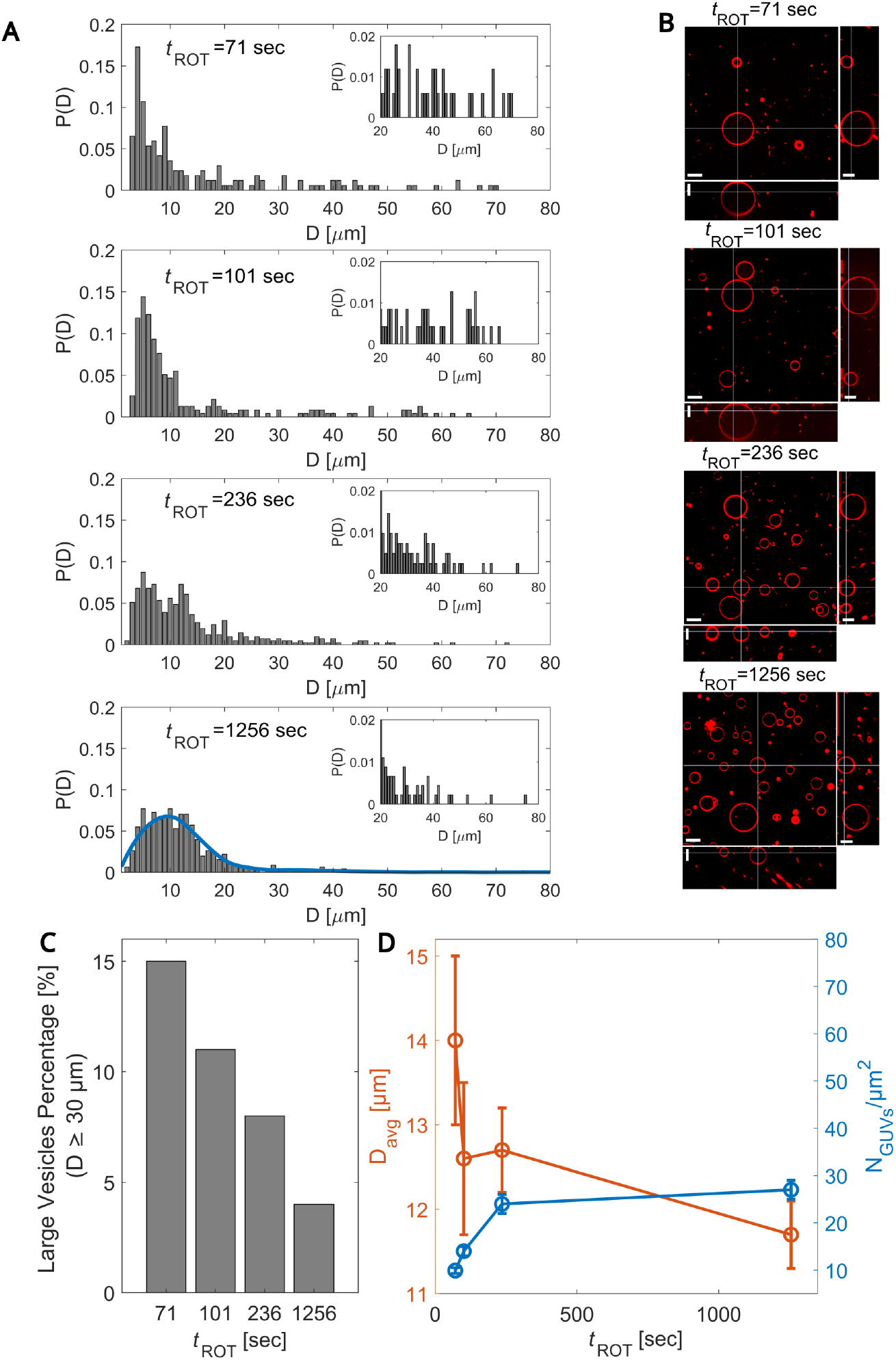
Effect of rotation time (*t*_ROT_) on GUVs’ size distribution. A) The panels show the probability distribution function P(D) for vesicle diameter D at a fixed angular frequency *ω* = 1600 rpm for four different rotation times = 71, 101, 236, and 1256 sec (top to bottom panels). Using quadratic regression, the blue curve is a fit to the base distribution, as measured from the longest experiment (*t*_ROT_ = 1256 sec). Insets: Zoom-in of the *P*(*D*) for *D* values ≥ 20*µ*m. B) Spinning disk confocal micrographs of GUVs in the xy cross-section (top view), the xz cross-section (bottom side view), and the yz cross-section (right side view) for the same running conditions as in (A). Images were corrected for index refraction mismatch between the objective immersion liquid, and our aqueous sample. ^20,21^ Scale bars are 20*µ*m. Membrane composition (molar percentage): 99.89% DOPC, 0.10% Liss Rhodamine-PE, 0.01% DSPE-PEG(2000)-Biotin. C) Percentage of large vesicles (*D* ≥ 30 *µ*m) *vs. t*_ROT_. D) GUVs diameter *D*_avg_ (orange) and GUVs density (blue) *vs. t*_ROT_. Data is presented as mean ± SEM. Number of vesicles used for data analysis: *N* = 168, *t*_ROT_ = 71 sec; *N* = 236, *t*_ROT_ = 101 sec; *N* = 412, *t*_ROT_ = 236 sec; *N* = 454, *t*_ROT_ = 1256 sec.

The histograms of the probability distribution *P*(*D*) for GUV diameters *D* at different *t*_ROT_ and spinning disk confocal micrographs of the generated GUVs for the same running conditions are depicted in Figs. 2A and B, respectively. Insets in Fig. 2A provide more detailed views of the larger size ranges. Our data show that as the rotational time decreases, the distribution shifts toward higher GUV diameters, with fewer observed GUVs (Figs. 2B and D) but a higher frequency of larger ones (Fig. 2A). These results are further supported by Fig. 2C which shows that the percentage of larger GUVs (*D* ≥ 30 *mu*m) is highest at the shortest rotational time of 71 sec and decreases monotonically as the rotation time increases. This shift in distribution supports the idea that shorter rotation times favor the arrival of large W/O droplets to the LOM/O interface. This size sorting effect leads to the observed increase in average diameter with decreasing rotation time, shown in Fig. 2D. However, this size sorting is negligible as the GUVs distribution does not change with a further increase in rotation time (not shown). As such, the distribution measured at long rotation time (1256 sec) provides a direct and quantitative measure of the input W/O droplet distribution introduced in the system, which we use as input for the theoretical model (see Theoretical model section below).

#### Effect of Angular Frequency (ω)

We now turn to explore the effect of varying the rotation frequency *ω* for a rotation time of 45 sec on size sorting. Fig. 3 depicts the data obtained for *ω* = 1200, 1600, and 2000 rpm and corresponding rotation times *t*_ROT_ of 97, 101, and 105 sec (including 40 sec dripping time and *ω*-dependent deceleration time). As in the effect of varying *t*_ROT_, we find that as *ω* decreases, the distribution shifts toward higher GUVs diameters with improved size sorting (Fig. 3A), with fewer observed GUVs (Figs. 3B and D) but a higher frequency of larger ones (Fig. 3A). These results are further supported by Fig. 3C, showing that the percentage of larger GUVs (*D* ≥ 30 *µ*m) is highest at the lowest *ω* = 1200 rpm and decreases monotonically as the rotation speed increases. As a result, the average GUV diameter also increases (Fig. 3D).

**Figure 3:**
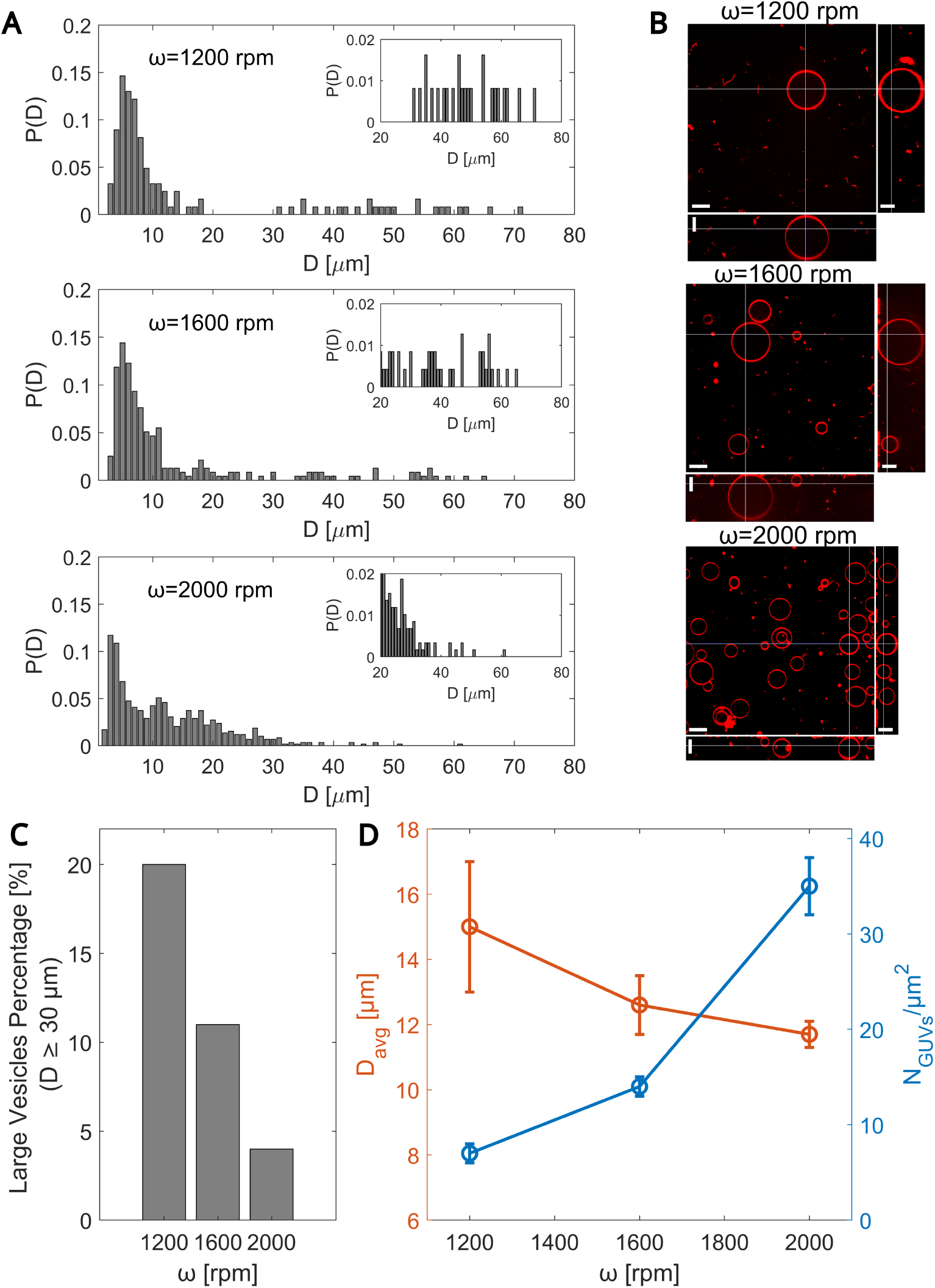
Effect of the angular frequency (*ω*) on GUVs’ size distribution. A) The panels show the probability distribution function P(D) for vesicle diameter D for variable *ω* values of 1200, 1600, and 2000 rpm (top to bottom). The rotation time *t*_ROT_ is 40 + 45 = 85 sec plus the deceleration time, yielding: 97, 101, and 105 sec, respectively. Insets: Zoom-in of P(D) for *D* ≥ 20*µ*m. B) Spinning disk confocal micrographs of GUVs in the xy cross-section (top view), the xz cross-section (bottom side view), and the yz crosssection (right side view) for the same running conditions as in (A). Images were corrected for the index of refraction mismatch between the objective immersion liquid and our aqueous sample. ^20,21^ Scale bars are 20*µ*m. Membrane composition as in Fig. 2B. C) Percentage of large vesicles (*D* ≥ 30*µ*m) *vs. ω*. D) GUVs diameter *D*_avg_ (orange) and GUVs density (blue) *vs. ω*. Data is presented as mean ± SEM. Number of vesicles used for data analysis: *N* = 123, *ω* = 1200 rpm; *N* = 236, *ω* = 1600 rpm; *N* = 589, *ω* = 2000 rpm.

Overall, we demonstrate that both *t*_ROT_ and *ω* can function as useful control parameters for optimizing the size sorting in the system. In particular, to increase the proportion of large vesicles in the sample.

### Theoretical Model

A model was developed to clarify the described phenomena by analyzing the balance of forces acting on a moving aqueous droplet within the cDICE chamber. The Weber number *We* quantifies the ratio between the stress exerted on the droplet by the surrounding oil phase, leading to distortion from a spherical shape, to the stress by surface tension, tending to preserve a perfectly spherical shape: ^22,23^

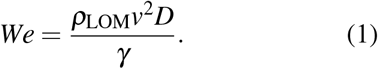

Here, *v* is the velocity of the droplet in the radial direction relative to the embedding medium, and *D* is the droplet’s diameter. As a crude estimate, we use *ρ*_LOM_ ≈ 1000 kg/m^3^, *σ* ≈ 50 mN/m, *D* ≈ 20 *µ*m and *v* ≈ 1 mm/s, and find that *We* ∼ 10^−6^. Since *We* ≪ 1, we conclude that droplets remain nearly spherical. ^24,25^

The acceleration 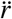 for a spherical droplet of radius *R* is given by

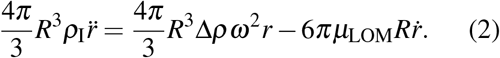

The first term on the right-hand side is the centrifugal force, while the second is the drag (Stokes) force acting on it.

The solution to this second-order linear ordinary equation is

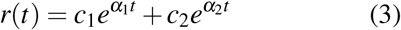

where the time constants *α*_*i*_ (*i* = 1, 2) are the roots of

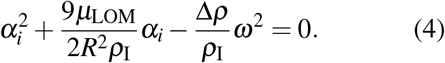

The constants *c*_1_ and *c*_2_ are fully determined from the initial conditions *r*(*t* = 0) = *r*_0_ and 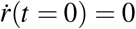, which translate to the two linear equations *c*_1_ + *c*_2_ = *r*_0_ and *α*_1_*c*_1_ + *α*_2_*c*_2_ = 0. The system has two inherent time-scales: *T*_1_ = 2*R*^2^*ρ*_I_*/*9*µ*_LOM_ and 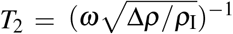. In the values of parameters involved, *T*_2_ ⪢ *T*_1_ meaning the left-hand side in Eq. 2 nearly vanishes, as originally proposed, ^18^ in agreement with the small Reynolds number *Re* ≈ 10^−3^ (viscous drag forces dominate the droplets’ motion).

Fig. 4 shows the theoretical predictions for three droplets with different radii. The smallest droplet (diameter 12 *µ*m) travels slower than the other two (diameters 16 *µ*m and 20 *µ*m). This is because the ratio between volume forces, scaling as ∼ *R*^3^, to surface forces (∼ *R*) is smaller than for the large droplets. In all the cases, the trajectories *r*(*t*) are almost linear on short times. In the simulation, we used a constant value of *ω* for a certain time followed by a linearly decreasing *ω*(*t*) as the experimental protocol prescribes (see Materials and methods for details). The curves *r*(*t*) level off at the end due to the deceleration period. Within the rotation time given here (97 sec), the droplets with diameters 16 *µ*m and 20 *µ*m traverse the entire LOM, while the smallest droplet does not. After, e.g., 40 sec, only the largest droplet (*D* = 20 *µ*m) reaches the interface and becomes a vesicle.

**Figure 4:**
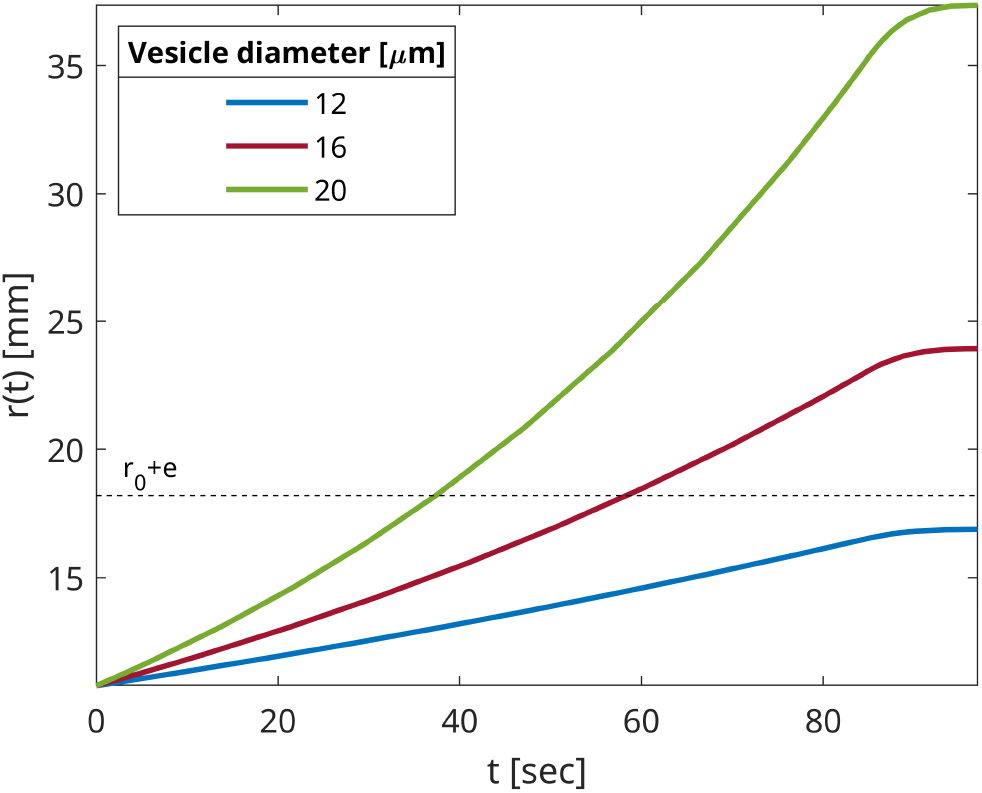
Three trajectories *r*(*t*) found from Eq. (2) for three droplet diameters at a rotation time of 97 sec. We used *ω* = 1200 rpm for a rotation time of 85 sec and *ω* linearly decreasing to zero at a rate of 100 cycles per sec in the final 12 sec. The horizontal dashed line is the outer radius, *r* = *r*_0_ + *e*. In this and in other model results, we used *r*_0_ = 10.8 mm, *e* = 7.4 mm, *µ*_LOM_ = 0.00564 Pas, *ρ*_I_ = 1040 kg/m^3^, and *ρ*_LOM_ = 898 kg/m^3^.

**Table 1:** Parameters Description.

| Parameter | Description |
| --- | --- |
| $\mu_{\text{LOM}}$ | Oil mixture viscosity |
| $\rho_{\text{LOM}}$ | Oil mixture density |
| $\rho_{\text{O}}$ | Outer solution density |
| $\rho_{\text{I}}$ | Inner solution density |
| $v$ | Droplet velocity |
| $R$ | Droplet radius |
| $r$ | Travel distance |
| $r_0$ | Distance of the oil layer from the rotation axis |
| $e$ | Oil layer thickness |
| $\omega$ | Angular frequency |
| $t_{\text{ROT}}$ | Rotation time |

One may also look at the effect of varying rotation time *t*_ROT_ at a fixed rotation speed *ω* = 1600 rpm. In Fig. 5, we calculated the GUV size distribution *P*(*D*), defined as the relative probability of finding a droplet with diameter *D*. In A, there are four panels for four times: 71, 101, 236, and 1256 sec. In each panel, the blue curve is a fit to the base distribution *P*(*D*) in the absence of rotation, taken from the experiments with the longest time of 1256 sec. The dashed vertical line marks the value of droplet diameter below which droplts cannot cross the LOM at the time of the experiment. The shaded region marks the droplet sizes that cross the LOM. The red curve is the predicted biased distribution of droplets *P*′(*D*) arriving at the outer radius of the device, *r*_0_ + *e*, at time *t*_ROT_ (Materials and Methods). It is clear that the shorter the experiment, the more *P*′(*D*) deviates from the base distribution *P*(*D*). Part C shows the average vesicle diameter *vs. t*_ROT_ based on theoretical predictions. Clearly, the average size decreases with time.

**Figure 5:**
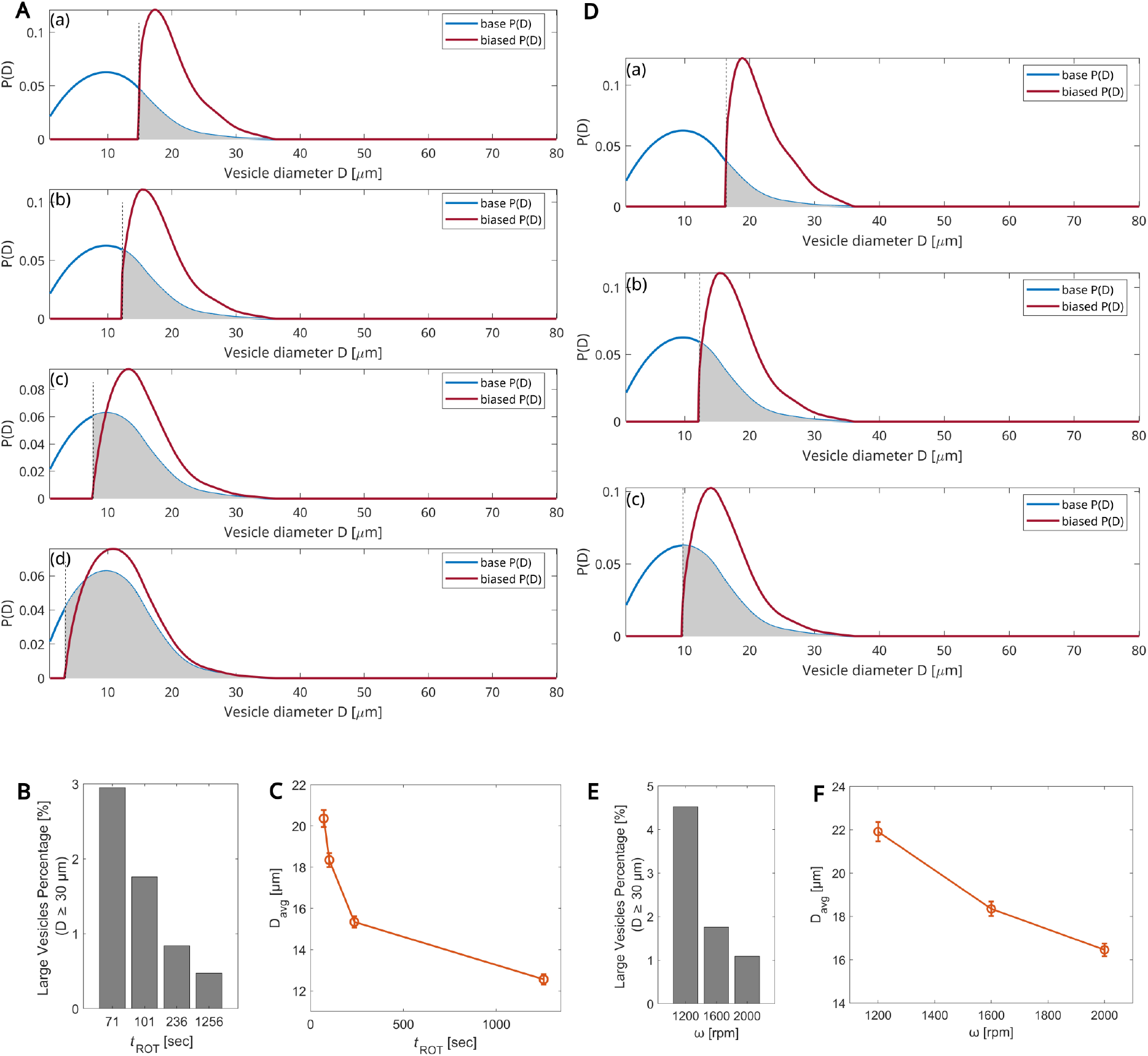
Predictions of the theoretical model. A) Effect of rotation time. Panels show the probability distribution function *P*(*D*) for vesicle size *D* at a fixed angular frequency *ω* = 1600 rpm for four different rotation times *t*_ROT_ = 71, 101, 236, and 1256 sec (top to bottom panels). The blue curve is a fit to the base distribution, as measured from the longest experiment (*t*_ROT_ = 1256 sec), using quadratic regression. The red curve is the calculated biased *P*′(*D*). B) Percentage of large vesicles (*D* ≥ 30 *µ*m) *vs. t*_ROT_. C) Mean diameter *vs. t*_ROT_. Data is presented as mean ± SEM. D) Predictions of the theoretical model for the effect of the angular frequency. *ω* varies between 1200, 1600, and 2000 rpm (top to bottom panels). The total rotation time is 40 + 45 = 85 sec plus the deceleration time. E) Percentage of large vesicles (*D ≥* 30 *µ*m) *vs. ω*. F) Mean diameter size *vs. ω*. Data is presented as mean *±* SEM.

In Fig. 5D, we simulated experiments with varying values of *ω* at a given time. Note that here, the experiment times are *t*_ROT_ = 97, 101, and 105 sec for the rotations speeds *ω* = 1200, 1600, and 2000 rpm, respectively. The three panels correspond to these three speeds. In each panel, the blue curve is the base distribution *P*(*D*) in the absence of rotation, taken from the experiments with the longest rotation time of 1256 sec. The red curve is the predicted biased distribution *P*′(*D*) of droplets arriving at the outer radius of the device at the given time *t*. The trend of decreasing mean diameter with *ω* is the same in both theory and experiment (Fig. 3D and Fig. 5F).

As a function of angular frequency or rotation time, the trends observed in both the theoretical model and experimental results consistently show that reducing angular frequency or rotation time increases the relative contribution of larger vesicles. This agreement suggests that the model may adequately explain the phenomenon. The model shows that at short rotation times and low angular frequencies, smaller droplets do not pass to the outer solution and, thus, do not become vesicles. However, probably due to other mechanisms, such as droplet-droplet interactions, smaller droplets still manage to pass the distance *r*_0_ + *e*, leading to deviations in the size distribution and the mean diameter values. Despite these differences, the overall trend remains consistent in theory and experiment.

## CONCLUSIONS

A modification of the cDICE method offers a relatively quick and straightforward, high encapsulation efficiency method for preparing GUVs for diverse research objectives. Our study demonstrates the technique’s effectiveness in sorting vesicles based on their size through simple systematic manipulation of key system parameters, including angular frequency and rotation time. The results of this approach provide valuable insights for researchers, allowing them to exert control over vesicle size. Multiple iterations can be conducted by utilizing the output as the new input to enhance control over the resultant sizes. This iterative process may aid in refining the requested sorting optimization. By providing a robust methodology for vesicle preparation and sorting, these findings are poised to significantly impact various fields. Notably, the ability to modulate vesicle size offers a considerable advantage for researchers developing versatile experimental systems, particularly in synthetic biology applications, as well as in drug delivery and bioreactor technologies.

## MATERIALS AND METHODS

### Materials

1,2-dioleoyl-sn-glycero-3-phosphocholine (DOPC, 850375C) and 1,2-distearoyl-sn-glycero-3-phosphoethanolamine-N-[biotinyl(polyethylene glycol)-2000] (ammonium salt) (DSPE-PEG(2000) Biotin, 880129C) were purchased from Avanti polar lipids in their solubilized form in CHCl_3_. 1,2-dioleoyl-sn-glycero-3-phosphoethanolamine-N-(lissamine rhodamine B sulfonyl) (ammonium salt) (Liss Rhodamine PE, 810150P) was purchased from Avanti Polar Lipids as a powder. Density gradient medium (Optiprep, D1556), Silicon oil (317667), Mineral oil (M5904), and Casein from bovine milk (C5890) were purchased from Sigma-Aldrich. D-(+)-Glucose, molecular biology reagent (02194024-CF) was purchased from MP Biomedicals, and *µ*-Slide 8 Well high (ibiTreat, 80826) was purchased from IBIDI.

## Methods

### The modified cDICE method for GUVs preparation

The various solutions and procedures are based on the protocols described in. ^6,15,19^ All steps are performed at room temperature (RT) unless stated otherwise.

### Lipid-in-Oil Mixture (LOM) Preparation

The LOM mixture is prepared following the protocol of. ^15^ Briefly, a lipid mixture with molar compositions of 99.89% DOPC, 0.10% Liss Rhodamine-PE, 0.01% DSPE-PEG(2000) Biotin and concentration of 6.65 mM was prepared in chloroform and stored under argon atmosphere at −20^°^C until used. In a subsequent step, an oil mixture consisting of 80% silicone oil and 20 % mineral oil (vol percent) was degassed for 2hr before being added to the lipid mixture to create a 0.417 mM LOM mixture with 6.25 vol% chloroform content. This mixture, which becomes turbid at this lipid concentration, ^15,17^ was kept on ice and used within a few minutes.

### Inner (‘I’) and Outer (‘O’) aqueous solutions preparation

Both solutions include glucose to control their osmolality, which is measured using an Osmometer (Gonotec Osmomat 3000). The inner solution is supplemented with 6.5% (vol) Optiprep and has an osmolality of 200 mosmol/kg, lower by 10 mosmol/kg compared to that of the outer solution.

### W/O emulsion preparation

This step is based on the protocol of. ^15^ Briefly, 700*µ*L of the LOM phase was added to 20*µ*L of the ‘I’ solution. To facilitate the formation of the W/O droplets, the solution was pipetted nine times with a 1000*µ*l pipette.

### Preparation of the GUVs via the modified cDICE method

Initially, 700*µ*L of the ‘O’ solution was introduced in the chamber, which rotates at a constant angular frequency *ω* (set to 1200, 1600, or 2000 rpm). In a subsequent step, 5 mL of the LOM was introduced in the rotating cDICE chamber. The preformed emulsion is then dripped into the chamber (center) using a 1000*µ*L pipette. This process takes typically 40 sec. Chamber rotation proceeds at the same constant speed for an additional period of 15, 45, 180, or 1200 sec, depending on the experiment, after which *ω* is gradually decreasing at a rate of 100 cycles/sec until halt. At this point, the system consists of two-separated, top (LOM) and bottom (GUVs), phases easily detectable when the chamber is tilted by 90^°^. The chamber is kept in this position for 10 min to promote GUVs’ sedimentation, after which 250 *µ*L of the bottom phase are collected and transferred in a casein-passivated Ibidi well. The wells are passivated with a solution of 2 mg/mL casein in 10 mM Tris-HCl (pH=7.5), incubated for 15 minutes, then rinsed first with 250*µ*L of DDW, and then with ‘O’ solution, and finally they are dried with a flow of *N*_2_. The same passivation procedure is employed for confocal imaging of the W/O emulsion.

### LOM viscosity measurement

The viscosity of the LOM was determined by a continuous rotation experiment. Measurements were carried out by MCR 702e MultiDrive rheometer (Anton Paar, Graz, Austria), equipped with stainless steel parallel plates geometry (*d* = 50 mm). The shear rate was conducted between 100 and 500 [1/s], and the viscosity [mPa sec] was recorded at 24^°^C.

### Microscopy technique

Imaging was performed using a Zeiss LSM 880 microscope in AiryScan mode or with a spinning disk confocal microscope equipped with a Yokogawa W1 module and a Prime 95B sCMOS camera (3i, Intelligent Imaging Innovations, Denver, USA). A 63X, .4 NA Corr.M27 Oil Immersion Plan-Apochromat objective was used for imaging. Images were corrected for index refraction mismatch between the objective immersion liquid, and our aqueous sample. ^20,21^

### Data quantification

We took confocal xy crosssection images (each of area 210*µ*m × 210*µ*m). The GUVs’ diameters were extracted manually by fitting their shape to a circle, from which their probability distribution function *P*(*D*) and mean diameter, *D*_*avg*_, was evaluated. Hundreds of vesicles were analyzed for each experimental condition (see Figure caption for details). To obtain a measure of the amount of GUVs that reached the ‘O’ phase, we estimated the number density of GUVs in units of *N*_GUVs_*/µm*^2^, averaged over 17 confocal images. Data quantification uses the software tools: Zen Black 2.1 (Zeiss, Germany) and ImageJ for raw data extraction and manual measurement of GUVs’ dimensions; MATLAB (MathWorks, MA, USA) for advanced analysis, data plotting, and curve fitting.

### Parameter values used for calculating the droplets traveled distance r

All parameter values are estimated at 25^°^C. *ρ*_LOM_ = 900 kg/m^3^, which we estimated from the volume average of the pure silicone (910 kg/m^3^) and mineral (840 kg/m^3^) oils density values (Source: Sigma-Aldrich website). Similarly, the density of the ‘I’ solution, *ρ*_I_ = 1040 kg/m^3^, was derived from the density values of the individual components (volume fraction of 0.065 OptiPrep (density =1320 kg/m^3^), 0.84 glucose 250 mM (density=1020 kg/m^3^), and 0.095 DDW (density=1000 kg/m^3^)). *µ*_LOM_ = 5.64 mPa sec, as determined experimentally. The cDICE chamber has dimensions of 38 mm in diameter and 7.4 mm in thickness. Combined with the volumes of LOM (5 mL) and the ‘O’ solution (700 µL) used, this results in *r*_0_ = 10.8 mm and *e* = 7.4 mm.

### Calculation of the biased vesicle size distribution P′(D)

*P*′(*D*) was calculated in the following steps. All vesicle diameters *D* were scanned; for a given *D*, the time *t*(*D*) for that particular size to cross the LOM was calculated based on Eq. (2). The time difference −*T* (*D*) was defined for *t*_ROT_ ≥ *t*(*D*) as −*T* (*D*) − *t*_ROT_ − *t*(*D*) and 0 otherwise. −*T* is proportional to the number of vesicles of size *D* arriving to the outer ‘O’ solution. The biased distribution is then obtained as *P*′(*D*) = *A* × −*T* (*D*)*P*(*D*), where the renormalization factor *A* ensures the sum of all probabilities is unity.

## Author Contributions

A.C. Performed experiments and data quantification, analyzed the experimental results, prepared the Figures, and wrote the manuscript.

S.G Developed the experimental system, designed experiments, and helped with data quantification.

L.O. Analyzed the experimental results.

E.Y., Constructed the rotating cDICE devices.

Y.T., Developed the theoretical model and wrote the manuscript.

A.B.G. Developed the experimental system, developed analytical methods for data quantification, and wrote the manuscript.

## Notes

The authors declare no competing financial interest.

## Data Availability

The data that support the findings of this study are available from the corresponding author upon reasonable request.

## ACKNOWLEDGMENTS

We thank Dr. Ran Tivony for valuable discussions and for reading the manuscript and Mattan Becker for assistance in constructing the cDICE device. We also thank Dr. Michal Zaiden for conducting viscosity measurements. A.B.G. is grateful to the Israeli Science Foundation (ISF) project no. 2101/20 and the Deutche Forschungsgemeinschaft (DFG, German Research foundation) project no. EL 1199/2-1 for financial support. A.C. gratefully acknowledges the Kreitman School of Advanced Graduate Studies for the Negev Scholarship. The authors gratefully acknowledge the IKI institute for Nanoscale Science and technology, Ben Gurion University of the Negev.

